# Spatio-temporal Reconstruction of Gene Expression Patterns in Developing Mice

**DOI:** 10.1101/2024.08.13.607727

**Authors:** Laura Aviñó-Esteban, James Sharpe

## Abstract

Understanding gene regulation in organism development is crucial in biology. Techniques like whole-mount *in situ* hybridization can reveal spatial gene expression in organs and tissues. However, capturing time-lapse images of gene expression dynamics in embryos developing *in utero*, such as mice, remains technically challenging beyond the early stages. To address this, we present a method to integrate static snapshots of gene expression patterns across limb developmental stages, creating a continuous 2D reconstruction of gene expression patterns over time. This method interpolates small tissue regions over time to create smooth temporal trajectories of gene expression. We successfully applied it to a number of key genes in limb development, including *Sox9*, *Hand2*, and *Bmp2*. This approach enables a detailed spatio-temporal mapping of gene expression, providing insights into developmental mechanisms. By estimating Gene Expression Patterns at previously unobserved time points, it facilitates the comparison of these patterns across samples. The reconstructed trajectories offer high-quality data that will be useful to guide computational modeling and machine learning, advancing the study of developmental biology in systems where real-time imaging is technically difficult or impossible.

## Introduction

The role of genes in regulating the development of a fully-formed functional organism is a central question in Biology. Morphogenesis can only be fully understood by characterizing the sequential unfolding of its dynamic processes, such as the regulatory interactions between genes and the patterns of gene expression over time [1, 2].

Gene expression patterns (GEPs) provide insights into the mechanisms of development. They can be visualized by labeling the target organ (or whole embryo) using techniques such as whole-mount *in situ* hybridization, which reveals which cells express specific genes. Genes that play crucial roles in organ development often exhibit distinctive patterns. For instance, in the developing limb bud, the gene *Hand2* is initially expressed in most posterior cells and has been shown to influence the posterior state of the limb bud [3] (Figure 1a). Similarly, *Sox9* is critical for cell differentiation into cartilage, and its expression dynamically evolves during limb development, gradually forming a pattern that mirrors the spatial arrangement of the skeleton [4] (Figure 1b).

**Figure 1:**
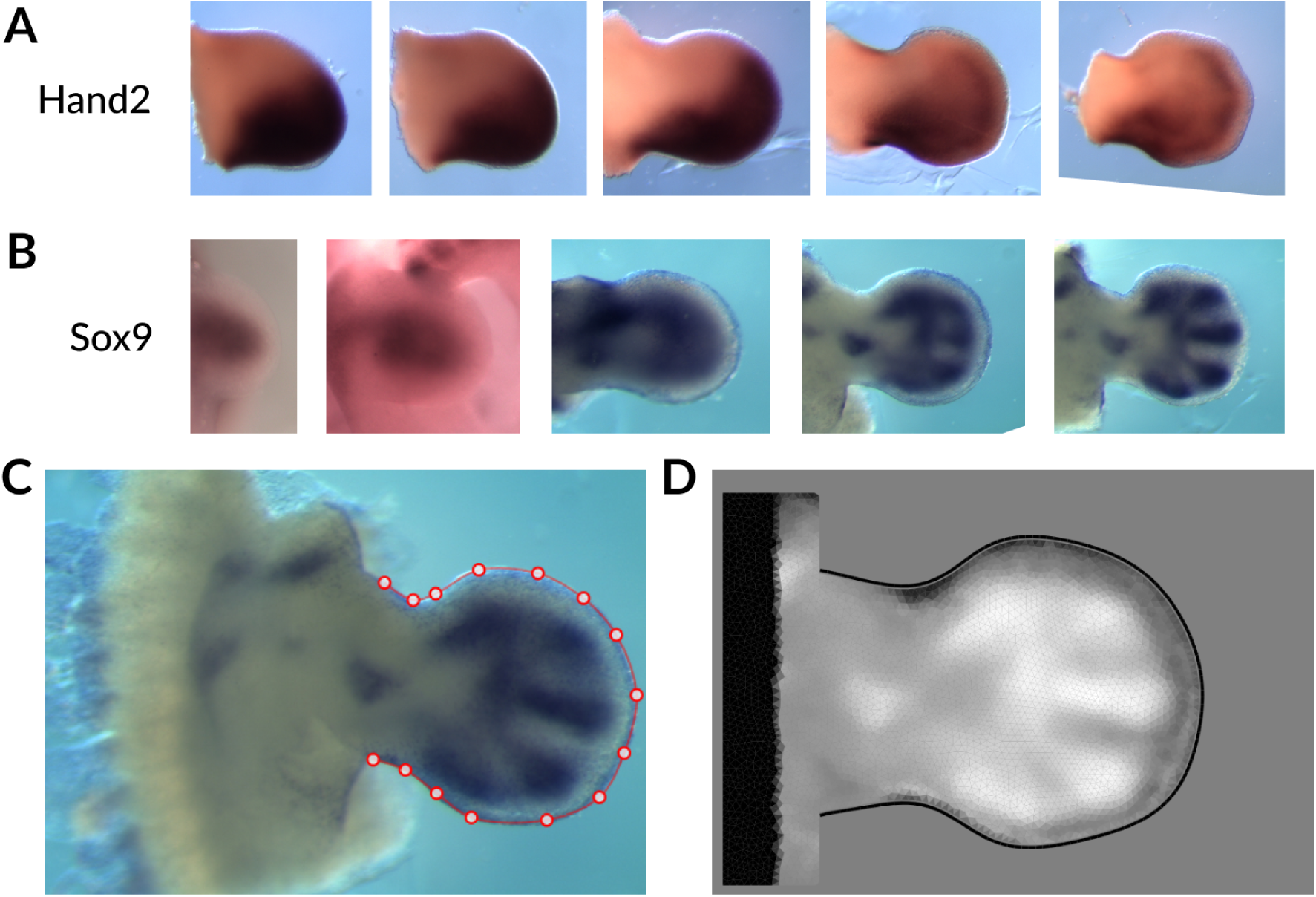
Gene expression patterns in the limb bud. (A) *In situ* Hyridzations of Hand2 gene. Developmental stage from left to right: E10:21, E11:01, E11:10, E11:18, E12:00. Note that the gene expression pattern in present on the anterior (bottom in this view) part of the limb. (B) *In situ* Hybridization of *Sox9* gene. Developmental stage from left to right: E10:11, E11:00, E11:12, E12:00, E12:12. The pattern of this gene converge to the skeletal pattern of the limb. (C) Staging method presented in [12, 13]. Red dots defines the outer shape of the limb (defined by the user). (D) Digitization of the gene expression pattern on digital meshes. White represents high expression levels, black represents low expression level.

Defining the boundaries of gene expression patterns (GEPs) presents a biological challenge. Firstly, gene expression can vary in intensity continuously across the domain. Consequently, delineating the spatial extent of gene expression can be ambiguous. Many genes in limb development exhibit “fuzzy” edges, with some showing clear gradient behaviors akin to morpho-gradients such as *FGF8* [5], while others display more subtle gradients. Secondly, many gene pattern boundaries do not align with anatomical boundaries. GEPs can dynamically shift through the tissue over time as cells regulate gene expression dynamically during development [6, 7].

Properly modeling of such a dynamical system requires a detailed description of the process. In externally developing organisms, such as Zebrafish or Drosophila, it is feasible to capture time-lapse movies of the entire developmental process *in vivo* in real-time [8, 9]. However, for internally developing embryos, this capability is currently limited beyond the gastrulation stage [10]. Moreover, *in vitro* culture techniques often fail to replicate full embryogenesis due to technical constraints. For example, in mouse, *in vitro* methods still cannot accurately mimic late embryonic stages beyond E10.5 [11].

One approach to address this challenge is by imaging fixed embryos, and their gene expression patterns, at different time points, gathering sufficient data to capture all dynamics. However, this approach relies on collecting a substantial amount of labelled image data, which may be time-consuming, and may also not be necessary for all genes. Additionally, creating such a dataset presents its own challenges. Firstly, biological noise is a significant factor. Even embryos from the same developmental time point may exhibit slight differences due to natural variation. Secondly, the rate of developmental progression can vary between embryos, even within the same litter. The developmental age of each embryo must be determined post-harvest using methods that estimate age based on observable features [12] (Figure 1c). Finally, a sufficient number of embryos must be harvested to ensure good coverage across different time points.

In this work we present a computational technique that allows us to construct a smooth, continuous description of GEPs over time and space with high temporal resolution. This is achieved by integrating static data samples collected from multiple individuals through computational tracking of moving tissue segments. To validate the efficacy of this method, we have applied it to key genes crucial for limb development and patterning, including *Sox9*, which is a marker of skeletal progenitor patterns cells. These reconstructed trajectories will provide a valuable reference for future modeling endeavors, significantly enhancing our ability to understand and predict developmental processes.

## Material and Methods

### Materials

In this study, we utilized a comprehensive dataset obtained from the Sharpe lab database, which includes *in situ*hybridization data collected over different projects. This dataset include a wide range of gene expression patterns in various developmental stages for both mouse hind limb and forelimb. The data were manually curated to ensure accuracy and consistency.

## Methods

### The Morphomovie and Tracking

We have previously named as a *Morphomovie* a sequence of meshes that represent a pre-defined growing 2D domain. (Figure 2a) This was first described in [14], and subsequently used in [15, 16, 13]. The morphomovie used in the current study describes the mouse hindlimb bud with a set of 292 meshes, corresponding to 48 hours of developmental time, with a 10 minute separation between each mesh. To transition from one mesh to the next one we just expand the edges of the elements to fit the new outer shape of the limb (Figure 2a). After each interval of 1 hour, a new mesh is created to maintain a consistent element size and account for the global tissue growth (Figure 2b). The internal tissue movements represented within our morphomovie approximate normal limb development, as these mesh movements were adjusted to fit real experimental data on labelled clones [14].

**Figure 2:**
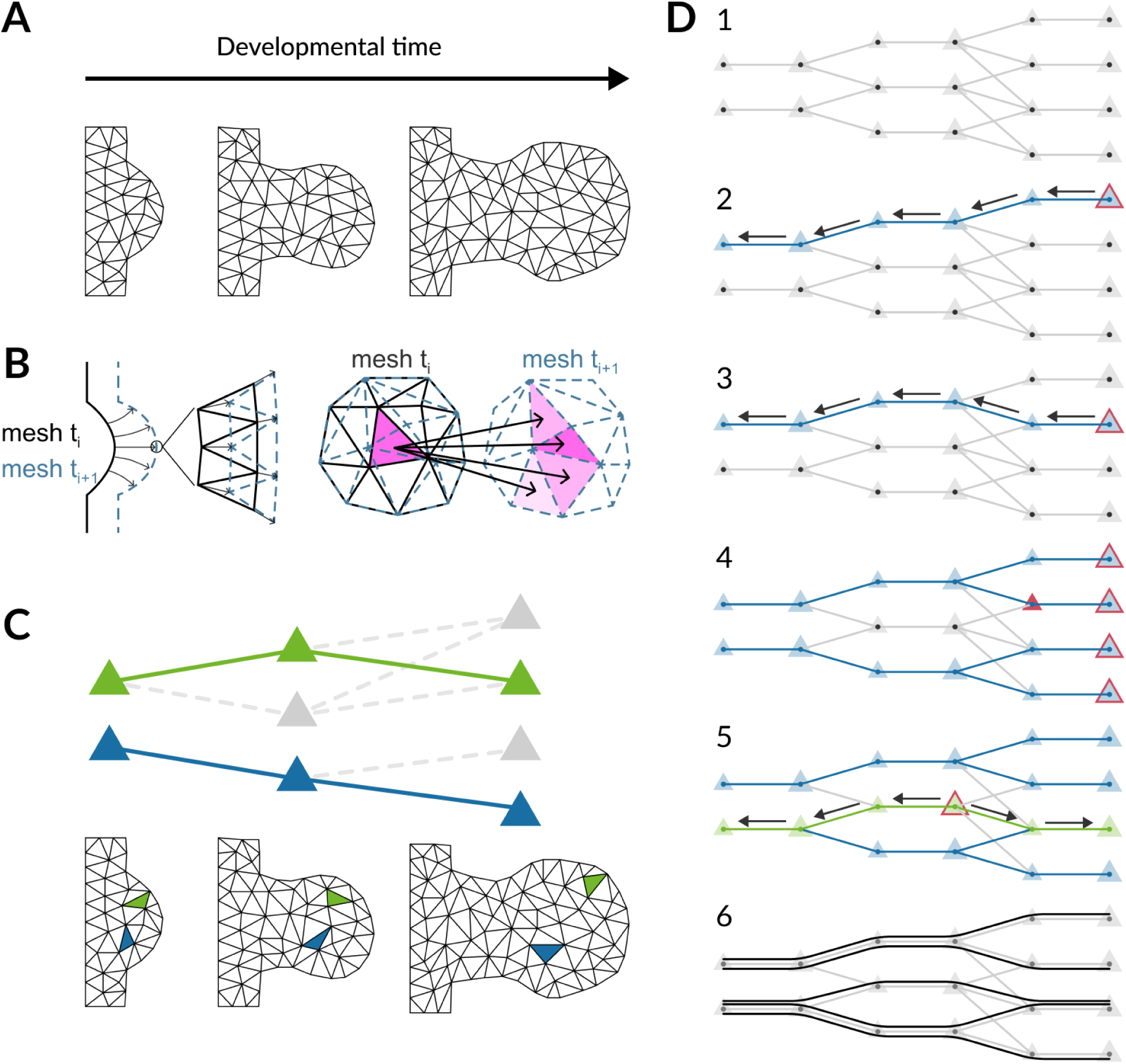
Tissue tracks. (A) Morphomovie toy example. The actual morphomovie has at higher resolution (i.e., each element has a smaller area). (B) Left: Every 10 developmental minutes, the mesh moves slightly to adjust to the growth and the new outer shape of the limb. The solid black line shows the outer contour at time t, while the dashed line shows the contour at time t+10 (in minutes). The triangles stretch to fill the new area. Right: Every 60 developmental minutes, there is an extra step of remeshing to maintain constant triangle size. The solid black line shows the outer contour at time t, and the dashed line shows the contour at time t+10 (in minutes). Each triangle can split into multiple triangles at the next time point, and one triangle can come from different triangles in the previous time point (adapted from [13]) (C) We can follow trajectories of triangles as a proxy for tissue movement in space. Top: Representation of the morphomovie as a Directed Acyclic Graph (DAG), showing predecessors and descendants of each triangle. (The 2 colors indicate 2 different example trajectories). Bottom: Spatial distribution of the trajectories. (D) Representation of the algorithm to derive the trajectories from the morphomovie. The size of the triangle illustrates growth. From top to bottom: (1) Representation of the Morphomovie as a DAG. (2) One triangle at the last time-point is picked and its trajectory tracked backwards by selecting the predecessors across developmental time. (3) This is repeated for the next triangle in the last time point. If one triangle has multiple predecessors (e.g., the red triangle), the algorithm selects the one with the higher area overlap as the predecessor. (4) This is repeated for all triangles at the last time point. (5) Some triangle at intermediate time-points are not picked in this process. For each of them a similar process as above is repeated, but this time both backwards and forwards so that all triangles are included in at least one trajectory, and all trajectories are complete from the first time-point to the last. (6) The final panel shows the different trajectories found in this toy example.

In our new interpolation technique, instead of dealing with the full 2D space at each timepoint, the space is divided into separate “tissue tracks” in time. We tracked the different tissue regions represented by triangles in the mesh. To achieve this we used the following approach. Due to the growth of the tissue, the final time point has more triangles than the initial time point. To ensure all triangles were captured in at least one trajectory, we tracked backwards from the last time-point to find the precursor triangle on the previous time point mesh. For the time points that were re-meshed, we picked the triangle with the highest area overlap. We performed this through all the time points for each of the triangles at the final time point (as shown in Figure 2c). It is important to notice, however, that because of remeshing, some triangles at intermediate time-points were not captured in any of this first set of trajectories. To correct for this problem, after each iteration of trajectory creation we search for triangles not yet captured and apply the same approach to them. However, since these triangles are not in the first or last time-point, we complete the “tissue track” backwards and forwards, so that all the trajectories are present in all time points. (Figure 2 c)

### Software used

In this study we used the staging system [12] and digitization through the web app LimbNET from the Sharpe Lab [13]. We use Vedo [17] to perform the B-Spline interpolation and the visualisation of the meshes. Other plots have been done using Matplotlib.

### B-spline Interpolation

Basis Splines, or B-splines, are a type of curves extensively used in computer graphics, animation, and computer-aided design. They create smooth interpolation curves by utilizing control points and specific basis functions, which manage the influence each point has on the curve. B-splines are designed to provide local control, meaning only nearby points affect the curve, unlike polynomial fits that can alter the entire curve with a single point adjustment. B-splines ensures smoothness of the curve an its derivatives. [18] This allow us to interpolate gene expression without abrupt changes in the expression or their rate of change.

### Histogram Matching

Given source histogram *h_s_* and reference histogram *h_r_*, the histogram matching process involves the following steps:

1. Compute the cumulative distribution function (CDF) of the source histogram *h_s_*: 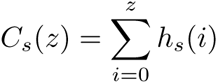
2. Compute the cumulative distribution function (CDF) of the reference histogram *h_r_*: 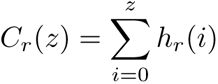
3. Create a mapping function *M* such that for each pixel value *z* in the source image, the corresponding value *z^′^* in the reference image is found using: 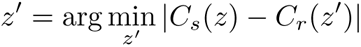

## Results

GEPs are spatial regions where a specific gene is expressed in the tissue. However, mapping GEPs from one time point to the next is challenging (see changing expression patterns in Figure 1). This difficulty arises because GEPs depend on the underlying cells and the movement of these cells over time. We therefore decided to adapt an interpolation approach focusing on how small tissue regions move over time. We broke the problem into many separate “tissue tracks” or trajectories instead of dealing with the full 2D pattern at each time point. This approach is a coarse-grained version of tracking individual cells movements and their gene expression changes over time. This is a reasonable approximation since genes in the limb mesenchyme do not have sharp on-off boundaries of expression. By calculating smooth temporal trajectories for how each small tissue region regulates gene levels (on, off, or intermediate), we can recombine all tracks in their correct spatial positions over time to recreate the full 2D GEP at any point. Consequently, we have transformed the interpolation task from a spatial problem to a largely temporal one.

### Tracking Tissue Trajectories

To represent the growing tissue, we used a collection of meshes called a morphomovie, a technique previously published in [14] and explained in methods. From the morphomovie, we extracted 42,155 “tissue tracks,” using triangles as the unit of tissue (Figure 2). These tracks are not entirely independent. Some trajectories overlap, but we consider this an advantage since, our goal is not analysis of tissue movements *per se*, but rather to create an average dynamic framework for interpolating GEPs. (Figure 2c).

### Intensity mapping and interpolation

After obtaining the tissue trajectories, we fitted the gene intensity data. We started with *Sox9*, a skeletal marker of the limb that shows high spatio-temporal complexity. The raw data includes 291 images from unstaged limbs. We used the staging system from [12] to semi-automatically assign a developmental age to each image. Then, to align the limb bud to the morphomovie, we morphed each image to the mesh corresponding to the assigned stage using [13]. This step estimates relative gene expression levels for all triangles at that time point. It is important to note that the experimental image data does not have uniform intensity levels across time. Each time point has a variable number of images, and some have none. This highlights the value of smooth interpolation over time. After the prepossessing of the images, the data includes 214 digitized images.

For each trajectory, we needed to smooth and infer the gene expression missing data using interpolation methods. A polynomial fit, which gives equal weight to all points, could be used, but it would struggle to capture subtle changes due to the constraints of the selected polynomial degree. Moreover, there is no prior polynomial (or curve) that the data should fit. As an alternative, piecewise linear interpolation could be used. To perform such interpolation, the mean intensity for each time point should be computed and then interpolated. This approach gives equal weight to each time point but not to each data point, and the resulting fit would not have any prior constraint. However, the result would not be continuous in the derivative, meaning a discontinuous rate of gene expression change, which we consider not biologically meaningful. Therefore, we chose to use B-splines, an interpolation technique that uses four neighboring points to define the sections, ensuring the final curve is continuous in the derivative by definition. We used Vedo [17] to perform the interpolation. Figure 3 a shows a graphical representation of the interpolation step. In 3b a real example trajectory is shown.

**Figure 3:**
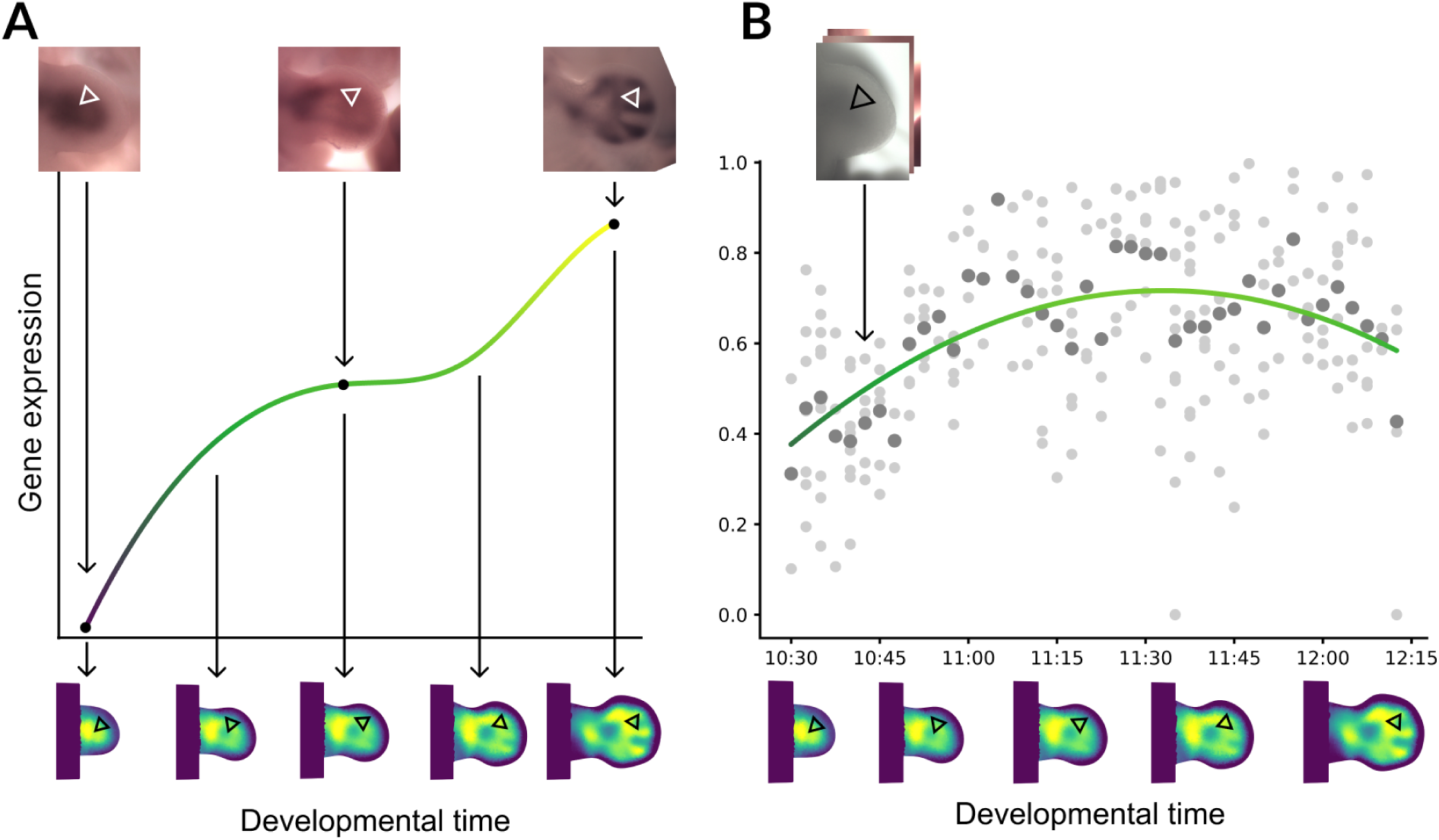
Gene expression interpolation. (A) Schematic of the process. Top row: Raw data for *Sox9 in situ* hybridization. White triangles represent a tissue track. Middle: Gene expression levels are obtained from the data of triangles in the tissue track. Then gene expression values are interpolated, represented by the line. Bottom row: The interpolation is map back to the morphomovie. (B) Real trajectory example. Light dots show the raw data. Dark dots show the mean value for each time point. The line shows the B-Spline interpolation.

Once each trajectory was interpolated, the values were mapped back onto the standard morphomovie. Triangles that were part of more than one trajectory are averaged. This provided a full 2D and time-continuous reconstruction of the *Sox9* gene (figure 4).

**Figure 4:**
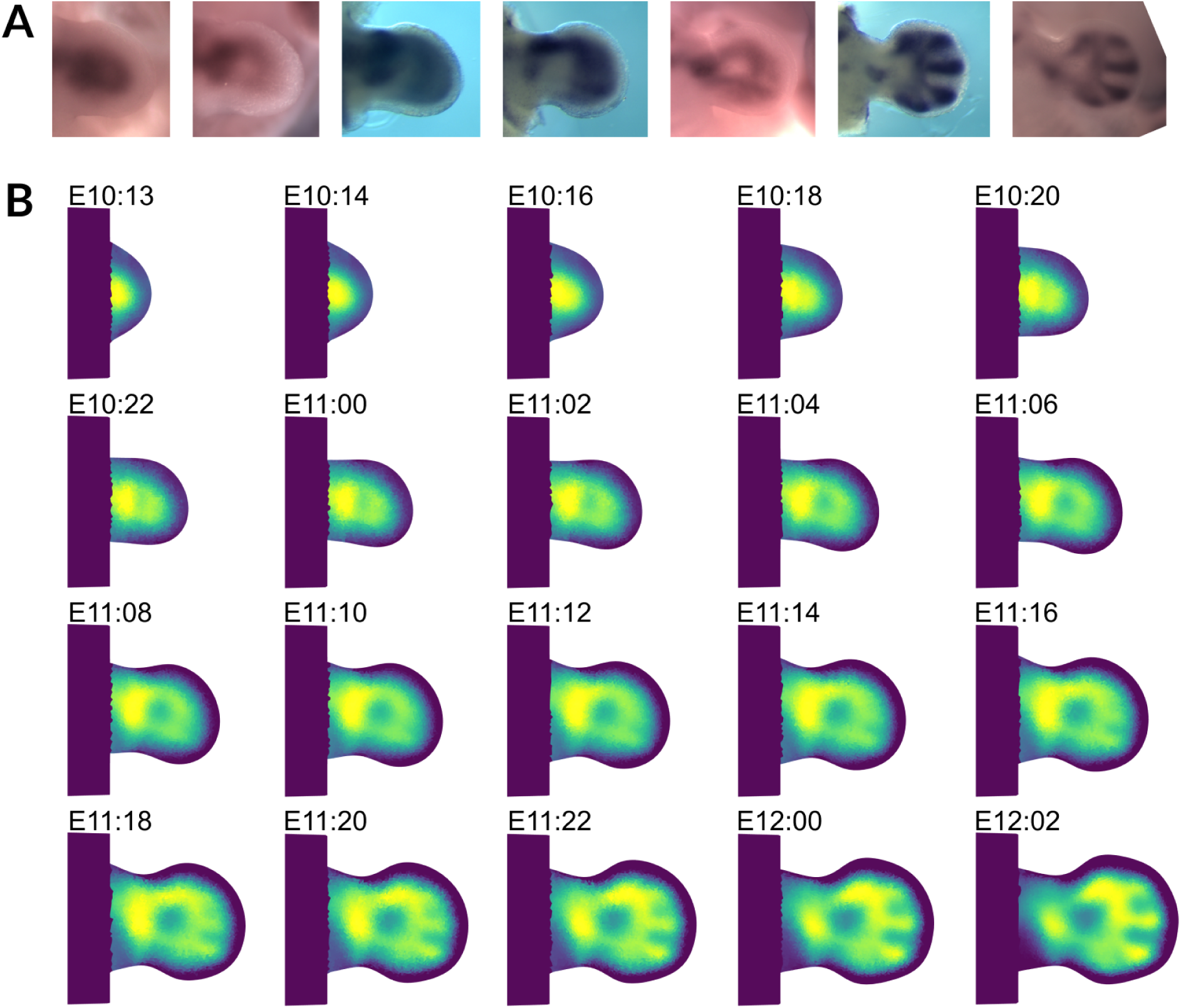
*Sox9* Reconstruction. (A) Subset of the *Sox9* raw data set. (B) Reconstruction of this gene from stage E10:13 to E12:02. Each reconstruction frame represents two developmental hours. Except first to second image that represents one developmental hour lapse.

### Refine the whole pattern using intensity optimization (boot-strapping)

One problem to address is that the raw data have inconsistent intensity levels from one image to the next due to experimental limitations and variation. In theory this should not be a major concern, as the information content we can obtain from *in situ* hybridization resides in the shapes of the GEPs rather than their absolute expression levels. Nevertheless, in practice it could reduce the quality of the result, and we therefore explored a boot-strapping approach to reduce this problem. To polish the overall pattern, we performed one cycle of bootstrapping, a refinement pass of the raw digitized data to reduce initial variation in expression intensity. Given that we cannot know an “absolute correct” set of expression levels for the data it is therefore non-obvious how to normalise the raw images *per se*. However, after estimating a first version of the GEP trajectory (previous section, and figure 4), this trajectory could be used as a template on which to normalise all the original raw images, and then recreate a new improved version of the trajectory from the new set of adjusted images. To perform normalization, we used histogram matching of the intensities of each raw image with the same time point from the first version of hte trajectory. Once all raw images had been adjusted, we used this new set to create a new reconstruction. Figure 5 graphically shows this procedure and the initial and final reconstructions for *Sox9* (Supplementary Video 1). As displayed in the figure, we managed to reduce initial noise, and also to remove some artifacts on the initial reconstruction.

**Figure 5:**
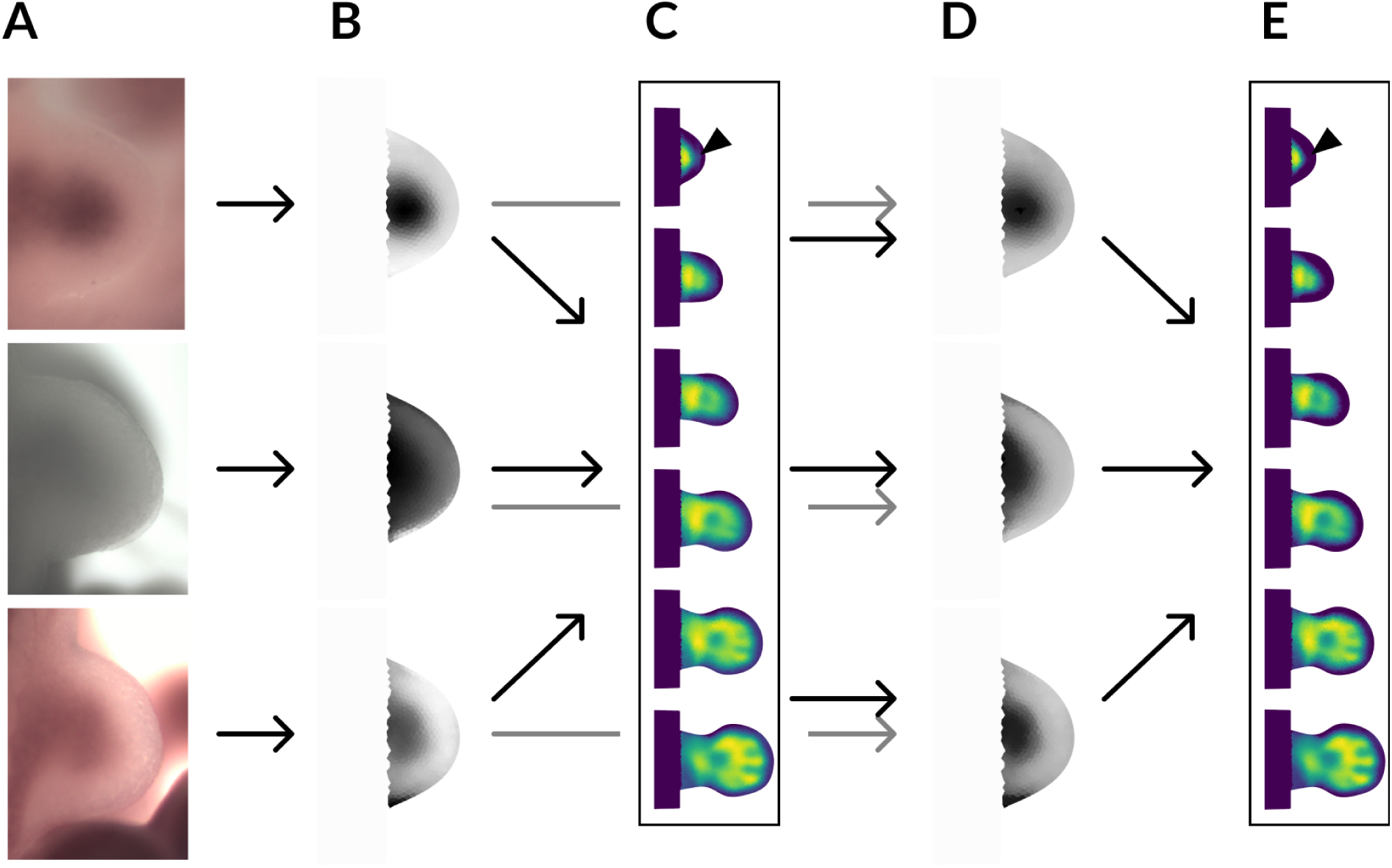
Refinement process for. (A) Raw data. Three images of Sox9 at the same time point E10:14, for which the variation in intensities is evident. (B) Digitization of the raw data. White represents no expression and black high expression. Note that the shape of the expression pattern is similar, yet the intensities show high variability. (C) Initial reconstruction using the digitization of the raw data. (D) Matching of the intensities of the raw data to the intensity values of the reconstruction at the same time point (E10:14) shows a highly improved digitisations compared to panel B. (E) Final reconstruction using the boot-strapped data. The process improves the quality of the reconstruction, for example correcting the artifact shown by the arrowhead.

### Application of the method to other genes

To showcase the generality of the approach, we generated 2D reconstructions of key developmental genes for the limb, including the anterior-posterior axis marker *Hand2* [19], a limb bud mesenchyme regulator *Twist1* [20], two chondrogenesis formation regulators *Bmp2* [21, 22] and *Wwp2* [23, 24], and an *FGF* downstream target *Dusp6* [25, 26]. We staged and digitised the images for these new genes. Then, we automatically applied the same algorithm that we used for *Sox9* to produce the reconstructions of each of these genes.

As shown in figure 6, we successfully managed to generate a smooth reconstruction for each of these genes, without any manual parameter tuning of the algorithm. Supplementary videos 2-6 show the reconstruction of these gene expression patterns for *Wwp2*, *Twist1*, *Bmp2*, *Dusp6*, and *Hand2*, respectively.

**Figure 6:**
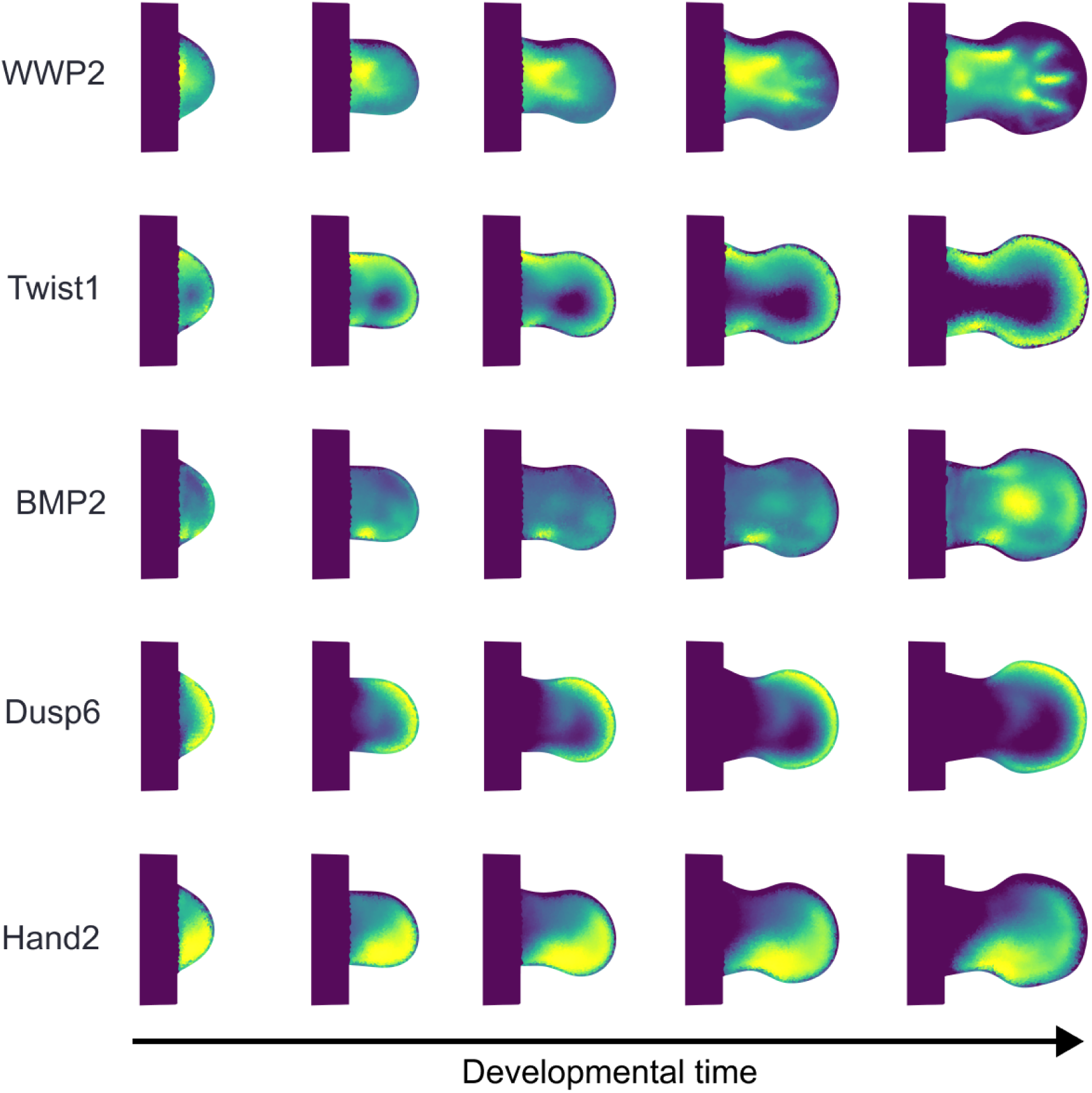
Application to other genes. From left to right the developmental stages E10:13, E10:22, E11:07, E11:16, E12:02. Note that the method has been able to capture simple patterns as well complex ones. Supplementary videos 2-6 show, respectively, the full reconstruction.

### Relationship between complexity of pattern and amount of image data required

An interesting question that arises from this approach is to asses or quantify how many images are needed to create a reasonable trajectory for a given gene. As seen in the previous sections, each gene has different spatial patterns over time. Some have complex shapes, while others have simple ones. It is therefore expected that the amount of data required to reconstruct the trajectory will vary: a simple pattern may not require many images, while a complex one will require a larger amount of data.

We explored this by picking two genes: *Dusp6* (simple pattern) and *Sox9* (complex pattern). We selected 10 images for each gene, evenly distributed over time, and applied our interpolation algorithm. 6 time-points of the interpolated trajectories are shown in 7A (Dusp6) and B (Sox9). Next, we chose a subset of just 3 of the raw images (the same inital and last images, and one in the middle of the sequence) and repeated the process. Figure 7 shows that with 10 images, we could reproduce the spatial dynamics of both *Sox9* and *Dusp6*. When reducing to just 3 input images, *Dusp6*’s trajectory was almost identical to the 10-image reconstruction. However, reducing *Sox9* to 3 images significantly impaired reconstruction quality, especially in the middle of the sequence. To go beyond visual inspection, we performed a quantitative comparison of the 2 reconstructions for each gene. The difference in pattern was calculated at each of the 6 time-points shown in 7, by summing the differences at every triangle in the mesh. Confirming what can be seen by eye, the difference in Sox9 pattern between the 10-image interpolation and the 3-image interpolation is greatest for the medium time-points (Figure 7C). In other words, when only 3 input images were used, the resulting interpolation diverges significant from the result using 10 images. The comparison for Dusp6 also shows an increase for medium time-points, but the overall differences are much lower. Since Dusp6 is a much simpler pattern fewer data-points can neverthless provide a good result. This suggests that in general complex patterns need more empirical data for accurate reconstruction.

**Figure 7:**
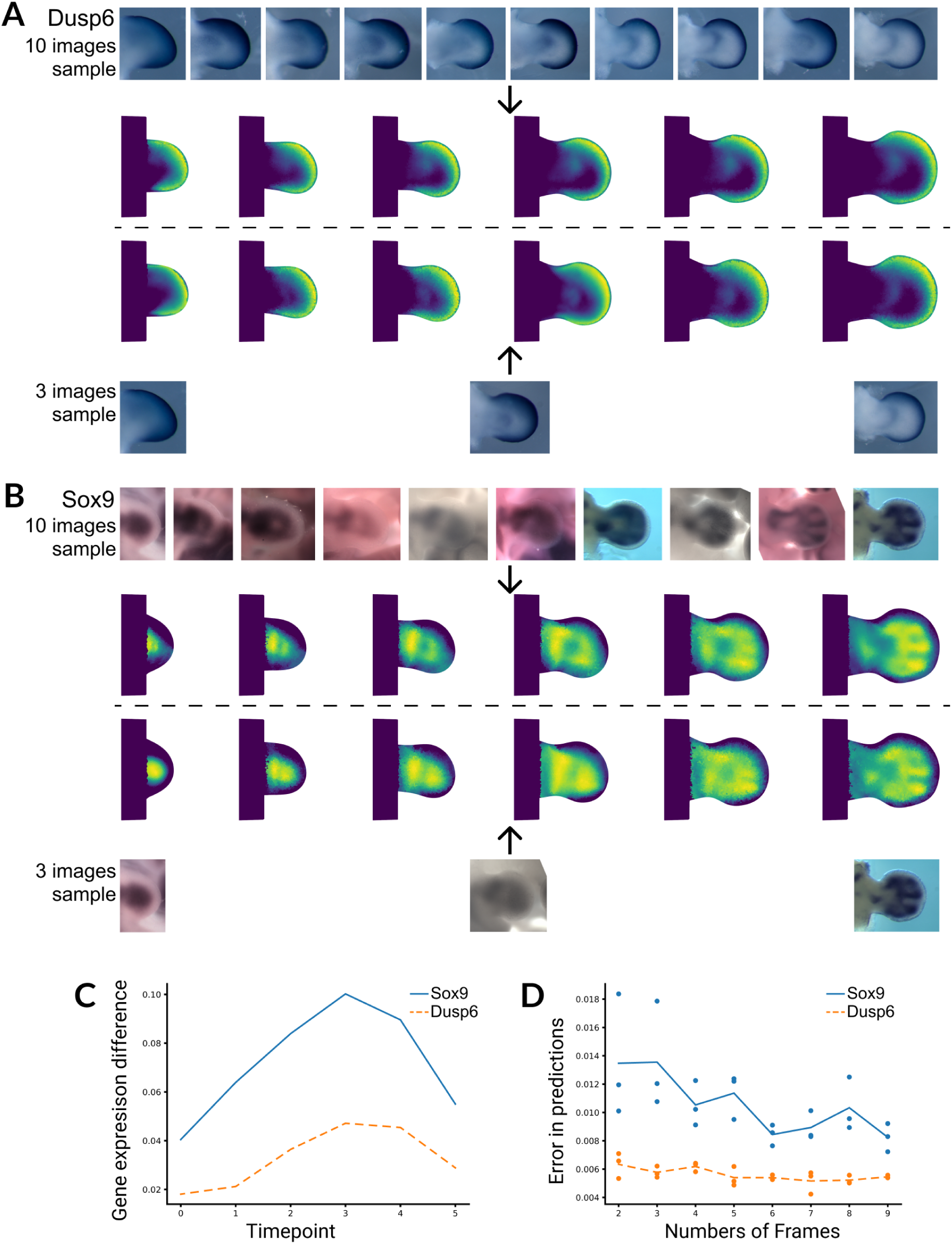
Reconstruction with different numbers of images. (A) *Dusp6* reconstruction. Top: The three selected raw images and their reconstruction. Bottom: The ten selected images and their reconstruction. (B) *Sox9* reconstruction. Top: The three selected images and their reconstruction. Bottom: The ten selected images and their reconstruction. (C) Comparing reconstructions of the same gene. We computed the residuals at each time point for reconstructions of the same gene. Solid Line: *Sox9* Residuals. Dashed Line: *Dusp6* Residuals. (D) The X axis shows the number of randomly picked frames used in the reconstructions. Each dot represents the residuals between five randomly selected images and the reconstruction.

To understand how the number of images affects the final reconstruction, we did a more comprehensive analysis - going beyond comparing just two amounts of input data (10 input images versus 3), and instead performing comparisons for the full range of the amounts of input data - from 4 input images to 10. We always selected the youngest and oldest image for each gene. To ensure each subset contained at least four unique time points, required for B-spline interpolation, we started with 2 intermediate random frames. For each subset, we compiled relevant data and performed a reconstruction. To calculate difference values, we needed to compare each predicted trajectory to some control trajectories. These control trajectories were calculated by randomly selecting five images from the original full dataset, excluding those used in the reconstruction. To reduce difference due to intensity profile variation of the raw data, we minimized the intensity difference between the reference and raw data by adjusting the raw data’s minimum and maximum intensity values. As shown in 7d *Dusp6* exhibits a flatter error line, indicating that its reconstruction is relatively stable regardless of the number of images used. In contrast, *Sox9* shows a significant decrease in errors as more data is added, highlighting the need of increased data for improving reconstruction accuracy for complex pattern genes. Additionally, for each particular number of frames used, *Sox9* displays higher variability in error measurements compared to *Dusp6*. This suggests that the reconstruction of complex shapes, as represented by *Sox9*, is less robust than that of simpler shapes like those of *Dusp6*.

## Discussion

A fascinating feature of gene expression patterns is that they do not necessarily correspond to fixed anatomical boundaries and can dynamically move through tissue during development. A specific cell might switch a gene on and later switch it off as the pattern “shifts” through that region of the tissue like a wave. In other words, the movements of tissue and GEPs do not always correlate.

In this study, we present a pipeline to reconstruct biologically meaningful continuous 2D and time spatiotemporal GEPs from discrete snapshots of expression over time in the developing mouse limb. The results directly provide a valuable insight into the temporal development of each gene. In particular, this tool can facilitate the comparison of correlations and differences in spatial patterns for different genes. Ideally, these comparisons should always be made at the same time point; our technique enables the estimation of a GEP at a time point where raw data may be unavailable. The final reconstructed trajectories can also be used as high-quality data to guide data-driven computational modeling and machine learning, as we have previously done with low-resolution reconstructions [16].

We have also provided a concrete demonstration of an intuitive idea - that the number of images needed for a good interpolation depends on the complexity of the GEP. Even though the number of images needed for a particular gene can not be predicted before hand, patterns which are highly dynamic, or spatially complex will need more images.

The approach is particularly useful for model systems for which it is impossible to capture time-lapse movies of gene dynamics and static images are used instead. This includes all later-stage mammalian embryos, and other species for which transgenic reporter constructs cannot be used. But as a tool, it may be useful for other species as well - helping to fill-in interpolated data from time-points which have not been captured. The main limitation of the method lies in the necessity of an independent staging, aligning and digitizing the original data to a standard reference. However, the tools provided here suggest that this method could indeed be applied to other model systems. In principle the same approach could also be generalized to 3D data by tracking tissue movements and interpolating intensity values for each track, providing that suitable 3D data exist.

## Acknowledgments

We thank Antoni Matyjaszkiewicz for invaluable discussions and help on use of the LimbNET API, Marco Musy for training on the use of vedo and the rest of the Sharpe Lab members. We also thank Giovanni Dalmasso for insightful feedback on the manuscript, and Maria Costanzo for her thorough manuscript review and constructive comments. The accumulated collection of whole-mount *in situ* data (from which images were taken for this paper: *Sox9*, *Wwp2*, *Twist1*, *Bmp2*, *Dusp6*, *Hand2*) was originally created by Jelena Raspopovic, Lucia Russo, Martina Niksic and Ju Yeon Han.

## Author Contributions

LA: Conceptualization, Data curation, Code Implementation, Formal analysis, Methodology, Writing – original draft; JS: Conceptualization, Supervision, Funding acquisition, Project administration, Writing – review and editing

## Supplementary Material

**Supplementary Video 1.** Interpolation of gene *Sox9* expression from developmental stage E10:12 to developmental stage E12:05. The video demonstrates the temporal progression with each frame spaced one developmental hour apart. A total of 214 digitized images were utilized to create this interpolation.

**Supplementary Video 2.** Interpolation of gene *WWP2* expression from developmental stage E10:21 to developmental stage E12:08. The video demonstrates the temporal progression with each frame spaced one developmental hour apart. A total of 15 digitized images were utilized to create this interpolation.

**Supplementary Video 3.** Interpolation of gene *Twist1* expression from developmental stage E10:17 to developmental stage E11:22. The video demonstrates the temporal progression with each frame spaced one developmental hour apart. A total of 17 digitized images were utilized to create this interpolation.

**Supplementary Video 4.** Interpolation of gene *BMP2* expression from developmental stage E10:14 to developmental stage E12:04. The video demonstrates the temporal progression with each frame spaced one developmental hour apart. A total of 16 digitized images were utilized to create this interpolation.

**Supplementary Video 5.** Interpolation of gene *Dusp6* expression from developmental stage E10:20 to developmental stage E12:04. The video demonstrates the temporal progression with each frame spaced one developmental hour apart. A total of 20 digitized images were utilized to create this interpolation.

**Supplementary Video 6.** Interpolation of gene *Hand2* expression from developmental stage E10:21 to developmental stage E12:00. The video demonstrates the temporal progression with each frame spaced one developmental hour apart. A total of 16 digitized images were utilized to create this interpolation.

